# Heterogeneity in VEGF Receptor-2 mobility and organization on the endothelial cell surface leads to diverse activation models by VEGF

**DOI:** 10.1101/800946

**Authors:** Bruno da Rocha-Azevedo, Sungsoo Lee, Aparajita Dasgupta, Anthony R. Vega, Luciana R. de Oliveira, Tae Kim, Mark Kittisopikul, Khuloud Jaqaman

## Abstract

The nanoscale organization of cell surface receptors plays an important role in signaling. We determined this organization and its relation to receptor activation for VEGF Receptor-2 (VEGFR-2), a critical receptor tyrosine kinase in endothelial cells (ECs), by combining live-cell single-molecule imaging of endogenous VEGFR-2 with rigorous computational analysis. We found that surface VEGFR-2 can be mobile or immobile/confined, and monomeric or non-monomeric, with a complex interplay between the two. The mobility and interaction heterogeneity of VEGFR-2 in the basal state led to heterogeneity in the sequence of steps leading to VEGFR-2 activation by VEGF. Specifically, we found that VEGF can bind to both monomeric and non-monomeric VEGFR-2, and, when binding to monomeric VEGFR-2, promotes dimer formation but only for immobile/confined receptors. Overall, our study highlights the dynamic and heterogeneous nature of cell surface receptor organization and its complex relationships with receptor activation and signaling.

## Introduction

There is growing evidence that the nanoscale organization of cell surface receptors plays an important role in their interactions with ligands and downstream effectors, and consequently cell signaling (Casaletto and McClatchey, 2012; Garcia-Parajo et al., 2014; Githaka et al., 2016; Jaqaman et al., 2011; Sungkaworn et al., 2017; Treanor et al., 2010; Zhou et al., 2017). This organization emerges from the complex regulation of receptor mobility, localization and interactions by various molecular and cellular factors. Therefore, it is imperative to study the dynamic organization of receptors and its relation to signaling in a receptor’s native cellular environment with minimal system perturbation. When compared to studies in reconstituted systems or model cell systems, studies in the native cellular environment reveal which behaviors indeed occur and how they relate to each other and to receptor function.

Live-cell single-molecule imaging is a powerful approach to observe individual molecules in their native cellular environment with high resolution in space and time (Liu et al., 2015; Xia et al., 2013). However, individual receptor behavior, be it localization, mobility or interactions, is generally heterogeneous and stochastic, especially under unperturbed conditions (Freeman et al., 2018; Jaqaman et al., 2011). In addition, individual receptor behavior and its related cellular-level processes span a wide range of spatial and temporal scales (Jaqaman et al., 2016). Therefore, computational and statistical analysis approaches are needed to tackle these challenges and attain a full understanding of the dynamic organization of cell surface receptors and its relation to receptor signaling.

Here we employed single-molecule imaging and a pipeline of automated image analysis and quantitative, multiscale data analysis to determine the dynamic organization of endogenous Vascular Endothelial Growth Factor (VEGF) Receptor-2 (VEGFR-2) in its native plasma membrane of endothelial cells (ECs), and link this organization to ligand binding and receptor activation. VEGFR-2, a receptor tyrosine kinase, is the main receptor in ECs for VEGF-A (commonly referred to as VEGF), the major promoter of angiogenesis in health and disease (Ferrara et al., 2003; Karaman et al., 2018; Olsson et al., 2006; Simons et al., 2016; Terman et al., 1992). Yet the mechanism of VEGFR-2 activation by VEGF is currently under debate. The classic model has been that VEGF binds to VEGFR-2 monomers, leading to their dimerization, activation by trans-autophosphorylation, and signaling (Ruch et al., 2007; Simons et al., 2016). In contrast, recent work in plasma membrane derived vesicles from CHO cells expressing exogenous VEGFR-2 provides evidence that VEGFR-2 can dimerize in the absence of ligand, such that VEGF binding to pre-existing dimers leads to dimer conformational changes that enable receptor trans-autophosphorylation and activation (Sarabipour et al., 2016). Our study allowed us to determine the interplay between these two competing models of VEGFR-2 activation, as part of providing a comprehensive and quantitative characterization of the hitherto unknown nanoscale spatiotemporal organization of VEGFR-2 on the EC surface.

## Results

### VEGFR-2 exhibits mobility and interaction heterogeneity on the surface of unstimulated ECs, with a complex interplay between the two

In order to image individual endogenous VEGFR-2 molecules in their native plasma membrane environment, we labeled early passage, unstimulated primary human microvascular ECs (pHMVECs) with a low concentration of primary Fab fragments (see Methods) that bound to the extracellular domain of VEGFR-2, followed by secondary Fab fragments conjugated to Rhodamine Red-X (RRX). The primary Fab fragments neither activated VEGFR-2 nor prevented its activation (Fig. S1A, B), thus allowing us to monitor endogenous surface VEGFR-2 with minimal perturbation. Total internal reflection fluorescence microscopy (TIRFM) was then used to acquire 10 Hz/20 s imaging streams (duration limited to 20 s to minimize photobleaching) at 37°C focused on the bottom surface of the cell (Fig. 1A; Movie 1), followed by multiple-particle detection and tracking (Jaqaman et al., 2008) to obtain the labeled receptor tracks (Movie 2).

**Figure 1.**
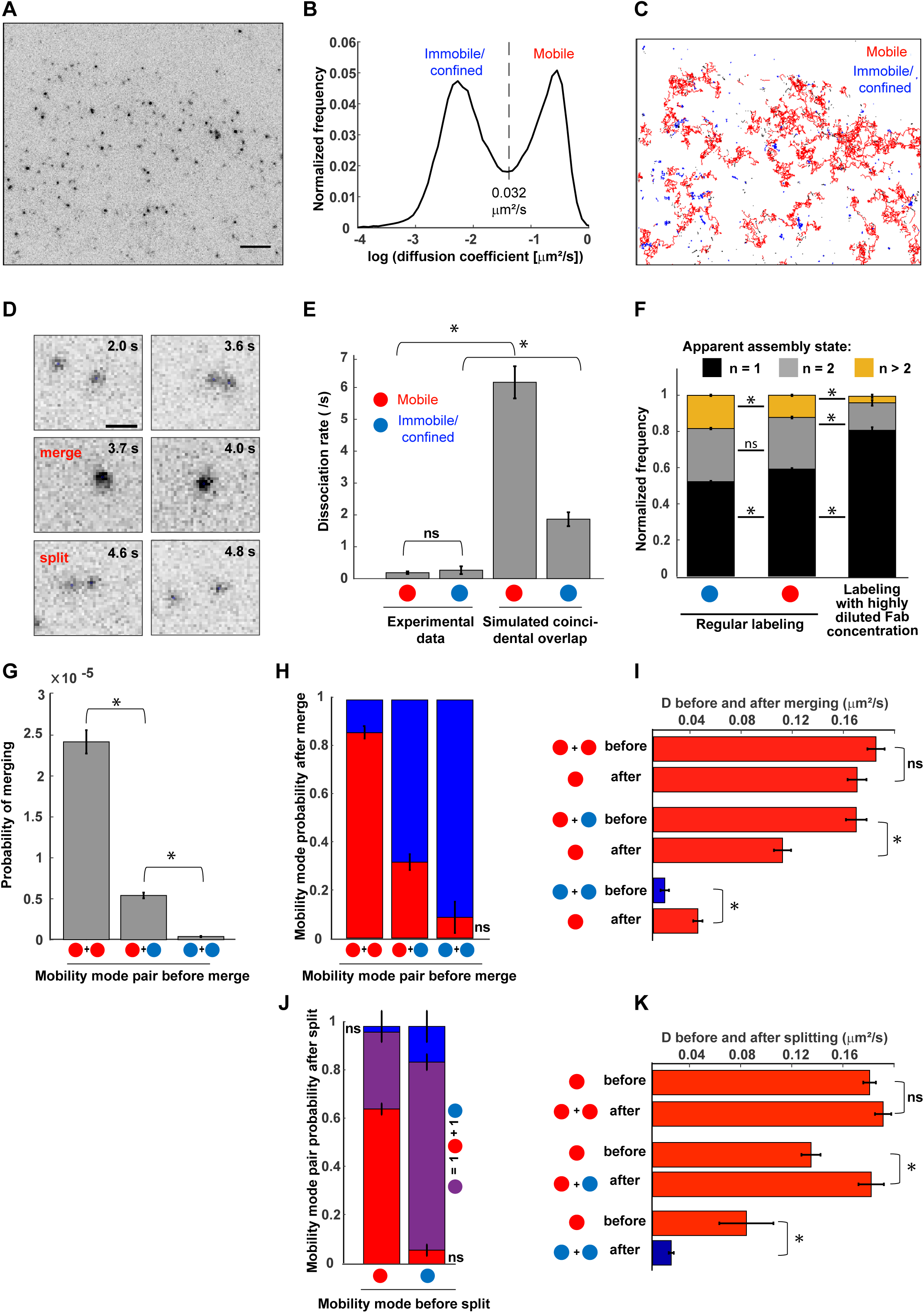
The dynamic organization of VEGFR-2 on the surface of unstimulated pHMVECs. All results are for unstimulated pHMVECs, so this will not be explicitly stated in individual panel descriptions. **(A)** Representative TIRF microscopy single-molecule image of RRX-labeled endogenous VEGFR-2 on the surface of a pHMVEC. Scale bar, 5 μm. Image inverted for visual clarity. **(B)** VEGFR-2 diffusion coefficient distribution. Vertical dashed line: threshold separating the mobile and imm/conf modes. **(C)** VEGFR-2 tracks over a 10 Hz/20 s imaging stream, colored based on mobility mode (red = mobile, blue = imm/conf, and black = too short to classify (duration < 5 frames)). Same cell and area as in (A). **(D)** Series of images showing a dynamic event of VEGFR-2 particle merging and splitting on the surface of a pHMVEC. Scale bar (applicable to all sub-panels), 1 μm. Images inverted for visual clarity. **(E)** Dissociation rate of VEGFR-2 per mobility mode as observed experimentally (“Experimental data”) and from simulations of coincidental overlap within the resolution limit (“Simulated coincidental overlap”). The latter was calculated by simulating non-interacting receptors that diffused in 2D in a manner similar to VEGFR-2 and then obtaining the distribution of apparent fusion times caused by resolution limitations (Jaqaman et al., 2011). Red and blue filled circles (also in F-K) represent mobile and imm/conf particles, respectively. **(F)** Frequency of apparent assembly state of detected VEGFR-2 particles in the two mobility modes and in samples with highly diluted labeling, the latter reflecting the number of fluorophores per secondary Fab fragment. **(G)** Probability of VEGFR-2 merging per combination of mobility modes. **(H)** Probability of mobility mode for merged particle based on the combination of particle mobility modes before merging. Red and blue bars (also in (I-K)) indicate mobile and imm/conf modes, respectively. **(I)** Comparison of VEGFR-2 diffusion coefficients before and after merging events that produce a mobile particle. If the two particles before merging have the same mobility mode (both mobile or both imm/conf), the displayed diffusion coefficient is the average of their two diffusion coefficients. If one particle is mobile and the other is imm/conf, then only the mobile particle diffusion coefficient is shown. **(J)** Probability of mobility mode combination for particles after splitting based on the mobility mode of the spitting particle. Purple bars indicate one mobile and one imm/conf particle. **(K)** Comparison of VEGFR-2 diffusion coefficients before and after splitting events that originate in a mobile particle. Analysis equivalent to that shown in (I), but for splitting instead of merging. Error bars in (E-K): standard deviation from 100 bootstrap samples. In (E-G, I, K), asterisks indicate p-value < 0.005 when comparing between the indicated conditions (assuming normal distributions with standard deviations obtained by bootstrapping); ns (not significant) indicates p-value > 0.05. In (H, J), ns (i.e. p-value > 0.05) indicates the probability is not significantly different from 0. N: VEGFR-2: 140,445 tracks (53,417 with duration ≥ 5 frames) from 9 independent experiments with 7-9 cells each; Highly diluted sample: 8751 tracks (2,200 with duration ≥ 5 frames) from 2 independent experiments with 10 cells each.

Diffusion analysis (Jaqaman et al., 2016) of individual VEGFR-2 tracks revealed a bimodal distribution of diffusion coefficients, allowing us to set a data-driven threshold (= 0.032 µm^2^/s) for VEGFR-2 surface mobility classification (Fig. 1B). Using this threshold, ∼35% of the tracks were classified as mobile, with a mean diffusion coefficient of 0.185 µm^2^/s, and the remaining ∼65% were much less mobile (mode termed immobile/confined (imm/conf)), with a mean diffusion coefficient of 0.008 µm^2^/s (Fig. 1C; Movie 3).

We also noticed heterogeneity in detected particle intensities (Fig. 1A), and merging and splitting events between detected particles (Fig. 1D; Movie 4), suggesting that VEGFR-2 was not in a purely monomeric state in unstimulated cells. These merging and splitting events largely reflected specific interactions between the imaged molecules, as the dissociation rate for VEGFR-2 interactions calculated from them (De Oliveira and Jaqaman, 2019) was in the range 0.19-0.26/s, which was about one order of magnitude lower than that expected from coincidental overlap due to diffusion (Fig. 1E). These specific interactions could reflect unligated dimerization or oligomerization (Bogdanovic et al., 2009; Chung et al., 2010; Lin et al., 2012; Low-Nam et al., 2011; Mischel et al., 2002; Sarabipour et al., 2016), indirect interactions through other receptors (Simons et al., 2016), or clustering within nanodomains such as clathrin-coated pits or caveolae (Labrecque et al., 2003; Lampugnani et al., 2006). To describe the non-monomeric state of VEGFR-2 in a neutral manner, we will use the term “assembly state.”

Estimation of the apparent assembly state *n* of the tracked VEGFR-2 particles from their intensities and sequence of merging and splitting events (De Oliveira and Jaqaman, 2019; Jaqaman et al., 2008) revealed a distribution of states, with more imm/conf particles in non-monomeric states than mobile particles (Fig. 1F). While a large fraction of the tracked particles had *n* = 1, confirming that VEGFR-2 labeling and imaging were at the single-molecule level, a substantial fraction had *n* > 1. Of note, because ∼20% of the secondary Fab fragments had two fluorophores (Fig. 1F, “labeling with highly diluted Fab concentration”), this analysis could not reveal the absolute assembly state of a detected particle. However, the fraction of VEGFR-2 with n > 1 was 41% for mobile particles and 48% for imm/conf particles, both significantly greater than 20% (p-value < 10^−15^; Fig. 1F).

To investigate further the relationship between VEGFR-2 mobility and interactions, we focused on the dynamic particle merging and splitting events (Fig. 1D; Movie 4). We found that mobile particles had a higher conditional probability of merging, especially with other mobile particles (Fig. 1G). At the same time, merging events enriched the imm/conf mode (Fig. 1H). This was consistent with the observation that imm/conf particles had an overall higher assembly state than mobile particles (Fig. 1F). For particles that stayed mobile after merging, there was some, albeit not always significant, reduction in their diffusion coefficient (Fig. 1I). Splitting exhibited the converse relationship, enriching the mobile mode (Fig. 1J, K). However, in contrast to the merging probability, the dissociation rate was the same for both mobility modes (Fig. 1E).

These analyses indicate that surface VEGFR-2 molecules in the plasma membrane of live, unstimulated pHMVECs exhibit heterogeneity in mobility and assembly state. The mobile mode of VEGFR-2 promotes interactions, which in turn enrich the imm/conf subpopulation, resulting in a complex interplay between VEGFR-2 mobility, interactions and assembly state.

### VEGF reduces VEGFR-2 mobility on the EC surface over multiple time scales

To determine how VEGF affects the dynamic nanoscale organization of VEGFR-2, shedding light on the role of this organization in VEGFR-2 signaling, we performed single-molecule imaging of VEGFR-2 after stimulating pHMVECs with 2 nM VEGF (specifically VEGF-A_165_), which led to robust VEGFR-2 activation that peaked at 5 min (Fig. S1C, D), as observed previously (Ferrara et al., 2003; Hamdollah Zadeh et al., 2008; Lampugnani et al., 2006; Simons et al., 2016). Because the effect of VEGF is time-dependent, and because the 20 s duration of a single-molecule imaging stream is too short to observe potential changes in VEGFR-2 behavior over minutes, we devised a “time-course” imaging and analysis strategy that allowed us to study single-molecule properties on time scales relevant to cellular processes. In brief, we imaged multiple cells within the course of an experiment (1 cell every ∼3 min, for ∼25 min total time), keeping track of their acquisition time relative to the addition of VEGF (usually just before imaging the second cell in a time-course; Fig. 2A). For data analysis (“time-course analysis” or TCA), multiple such time-courses were aligned either by the time point of VEGF addition, or time 0 (i.e. first cell) in the case of unstimulated cells. This allowed us to group cells based on time (∼15 cells per 5-min time interval, with 1000-1500 VEGFR-2 tracks each) in order to investigate their single-molecule properties over longer time scales than captured by any individual single-molecule imaging stream.

**Figure 2.**
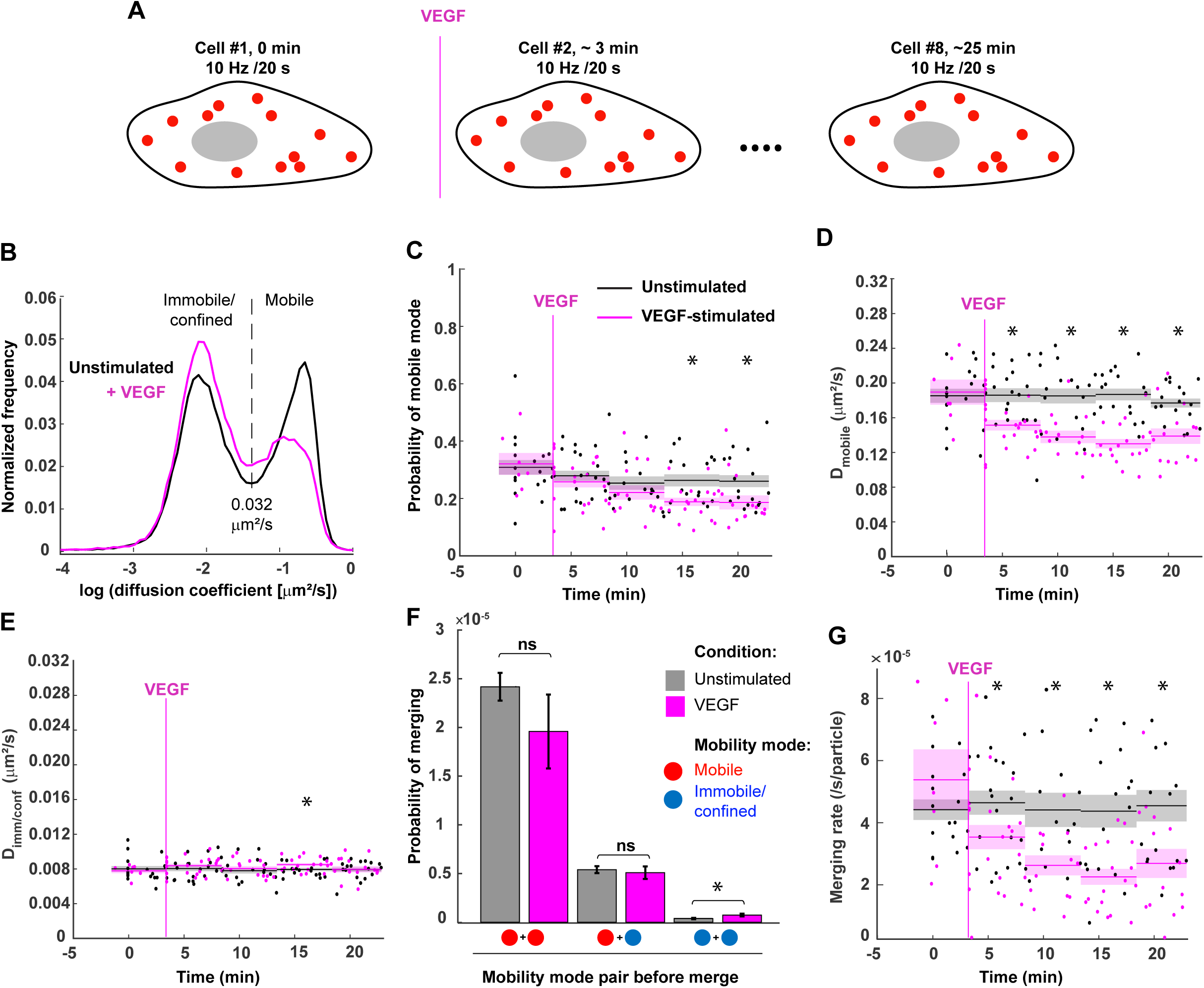
VEGF-induced changes in the dynamic organization of VEGFR-2 on the pHMVEC surface. **(A)** Cartoon illustrating time-course imaging. **(B)** VEGFR-2 diffusion coefficient distribution on the surface of unstimulated (black) and 2nM VEGF-stimulated (magenta) pHMVECs. Unstimulated distribution and vertical dashed line are a repeat of Fig. 1B. **(C-E)** Time-courses of the probability of VEGFR-2 molecules belonging to the mobile mode **(C)**, and the mean diffusion coefficient of mobile molecules **(D)** and imm/conf molecules **(E)** (note the different y-axis ranges in (D) and (E)). Black and magenta: measurements from unstimulated and VEGF-stimulated time-courses, respectively. Vertical magenta line: mean time point of VEGF addition (thus measurements at times before the vertical magenta line are from unstimulated cells, even in the stimulated time-course). Dots: individual cell values. Lines and surrounding shaded areas: the mean and standard error of the mean, respectively, from cells grouped into 5-min intervals as indicated. Asterisks above time intervals: p-value < 0.05 for comparing stimulated and unstimulated measurements per time interval using a t-test. Time intervals without an asterisk: no significant difference between stimulated and unstimulated measurements (p-value > 0.05). **(F)** Probability of VEGFR-2 merging per combination of mobility modes before the merge in cells stimulated (magenta) or not (gray) with 2 nM VEGF for 10-20 min. Error bars: standard deviations from 100 bootstrap samples. Asterisks and ns (not significant): p-value < 0.05 and p-value > 0.05, respectively, when comparing between the indicated conditions (assuming normal distributions with standard deviations obtained by bootstrapping). **(G)** Time-course of the rate of merging. All details as in (C-E). N: Unstimulated: same as in Fig. 1; Stimulated: 99,699 tracks (42,865 with duration ≥ 5 frames) from 8 independent experiments with 7-8 cells each.

Focusing first on mobility, our analysis revealed that VEGF led to a gradual decrease over time in the probability of the mobile mode of VEGFR-2, starting at 5-10 min after VEGF addition (Fig. 2B, C). Also, but in contrast to the gradual shift in population from mobile to imm/conf, the diffusion coefficient of mobile molecules (D_mobile_) exhibited an almost immediate drop upon VEGF addition, down to ∼0.135 µm^2^/s (Fig. 2D). The diffusion coefficient of imm/conf molecues (D_imm/conf_) did not change (Fig. 2E). These results indicate that VEGF has both a fast effect on VEGFR-2 mobility, where it reduces D_mobile_, and a slow effect, where it shifts the population of VEGFR-2 molecules toward the imm/conf mode. The slow effect occurs on the same timescale as the cellular signaling response (Fig. S1C, D), while the fast effect is almost immediate.

### VEGF enhances the probability of interactions between imm/conf VEGFR-2 molecules

Next we investigated the effect of VEGF stimulation on VEGFR-2 merging events, to determine whether VEGF promotes VEGFR-2 interactions, as described by the classic model of VEGFR-2 activation (Ruch et al., 2007; Simons et al., 2016). Interestingly, we found that VEGF stimulation increased the conditional probability of merging for imm/conf-imm/conf particle pairs, but not for events involving mobile particles (Fig. 2F). However, in spite of this increase in merging probability for imm/conf-imm/conf particle pairs, the overall rate of merging between VEGFR-2 particles decreased over time in the presence of VEGF (Fig. 2G). This rate was for all particles regardless of their mobility mode, akin to an overall association rate (but for only the labeled subset of VEGFR-2 molecules). This reduction in merging rate upon VEGF addition reflected the fact that most merging events involved mobile particles (as reflected by their much higher merging probability (Fig. 2F)), the fraction and diffusion coefficient of which decreased in the presence of VEGF (Fig. 2C, D).

These results suggest that the canonical model applies to interactions between imm/conf VEGFR-2 molecules, most likely because imm/conf receptors are limited to sampling a very small area of the cell surface, if any at all, and thus have a very low chance of encountering other imm/conf receptors on their own. In this case, VEGF, which is a constitutive dimer (Ferrara et al., 2003), can act as an attractor that brings receptors together. In contrast, mobile receptors have a much higher chance of encountering other receptors on their own, and VEGF does not seem to increase their interaction probability further.

### VEGF binds to both VEGFR-2 monomers and pre-existing non-monomers on the EC surface

The above analysis provides evidence that the canonical model of VEGFR-2 activation applies to a subset of VEGF-VEGFR-2 binding events on the surface of pHMVECs. To investigate to what extent the alternative model of VEGF binding to pre-existing VEGFR-2 dimers (Sarabipour et al., 2016) applies I the native cellular context, we performed simultaneous 2-color TIRFM imaging of VEGFR-2 (labeled as above) and VEGF (conjugated with Atto488, which did not interfere with VEGF function (Fig. S1E, F)). To increase the chance of observing colocalization between a labeled VEGFR-2 molecule and a labeled VEGF molecule, we labeled VEGF at a relatively high density (40% of the 2 nM dose was labeled). At this density, we were still able to detect single-particles of VEGF using Gaussian mixture-model fitting (Jaqaman et al., 2008). However, the density was sometimes too high for reliable tracking of labeled VEGF particles. Therefore, we tracked VEGFR-2 particles as above, and then associated VEGFR-2 tracks in each frame with co-localized VEGF particle detections (Fig. 3A; Movie 5). With this, we analyzed the interactions, assembly state and mobility of VEGFR-2 in the context of its association with VEGF, within the framework of TCA.

**Figure 3.**
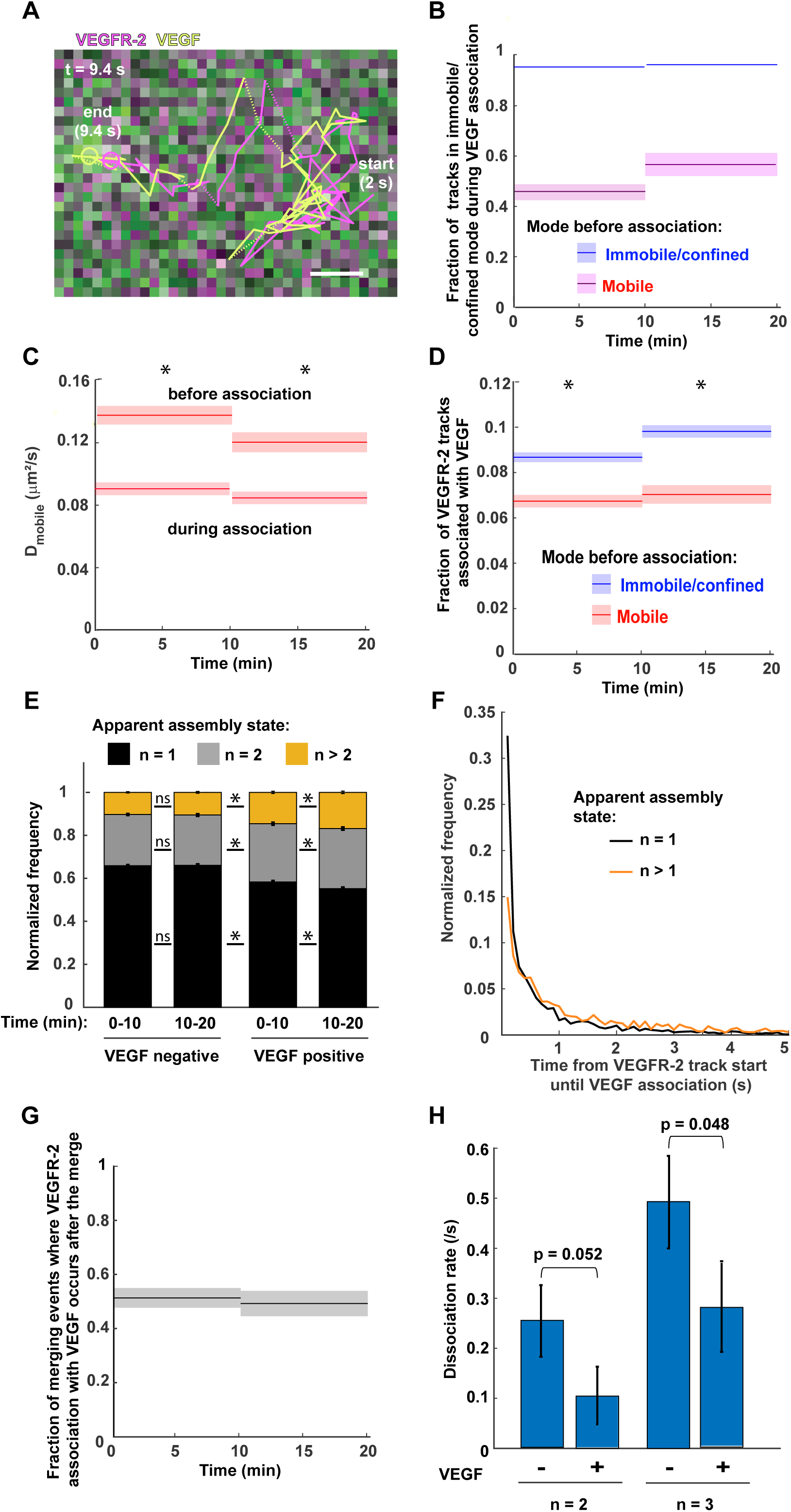
Dissection of the relationship between individual VEGFR-2 molecules associating with VEGF and their mobility and interactions. **(A)** A track of RRX-labeled VEGFR-2 (magenta) and its associated Atto488-labeled VEGF (light green), overlaid on the merged image of the 561 (magenta) and 488 (green) channels in the last displayed frame (at 9.4 s). Scale bar, 500 nm. **(B)** Time-course of fraction of VEGFR-2 tracks in the imm/conf mode while associated with an Atto488-VEGF particle, per mobility mode before VEGF association. Lines and surrounding shaded areas: measured parameter and its standard deviation from 100 bootstrap samples. **(C)** Time course of the diffusion coefficient, before and during VEGF association, of VEGFR-2 particles that are mobile both before and during VEGF association. Lines and surrounding shaded areas as in (B). Asterisks above time intervals: p-value < 10^−5^ for comparing the two measurements in each time interval (assuming normal distributions with standard deviations from bootstrapping). **(D)** Time course of fraction of VEGFR-2 tracks getting associated with Atto488-VEGF per VEGFR-2 mobility mode. Lines and surrounding shaded areas as in (B). Asterisks as in (C). **(E)** Frequency of apparent assembly state of detected VEGFR-2 particles associated (VEGF positive) or not (VEGF negative) with Atto488-VEGF, in the indicated time intervals. Error bars: standard deviations from 100 bootstrap samples. Asterisks and ns: p-value < 0.005 and p-value > 0.05, respectively, when comparing corresponding frequencies between the indicated conditions (assuming normal distributions with standard deviations obtained by bootstrapping). **(F)** Normalized frequency of waiting times between the start of a VEGFR-2 track and its association with Atto488-VEGF, for the subset of tracks that associate with VEGF after their first time point (23% of n = 1 tracks and 60% of n > 1 tracks). **(G)** Time-course of fraction of merging events where VEGFR-2 association with Atto488-VEGF occurs after the merge. Lines and surrounding shaded areas as in (B). **(H)** Dissociation rates of VEGFR-2 undergoing interactions associated (+) or not (-) with Atto488-VEGF, for n = 2 or n = 3 assembly states. Error bars as in (E). N: 189,097 tracks (81188 with duration ≥ 5 frames) from 10 independent experiments with 6-8 cells each.

Consistent with the global mobility trends observed above (Fig. 2C, D), this analysis revealed that, at the level of individual VEGFR-2 molecules, VEGF association both promoted the switch from mobile to imm/conf (Fig. 3B), and the reduction in D_mobile_ for receptors that stayed mobile while associated with VEGF (Fig. 3C). Interestingly, we found that a larger fraction of imm/conf VEGFR-2 particles got associated with VEGF than mobile VEGFR-2 particles, implying that VEGF had a higher probability to bind to VEGFR-2 already in the imm/conf mode (Fig. 3D). We reasoned that this could be because of the overall higher assembly state of imm/conf particles (Fig. 1F), which could increase the avidity of VEGF-VEGFR-2 binding (Fuh et al., 1998; King and Hristova, 2019).

The assembly state of VEGFR-2 particles positive for labeled VEGF was indeed higher than that of VEGFR-2 particles negative for labeled VEGF, and it increased over time (Fig. 3E). This analysis could however not inform us whether VEGF bound to monomers that then associated with each other, or whether it bound to pre-existing non-monomers. To distinguish between these two scenarios, we measured the waiting time from the start of each VEGFR-2 track to its moment of labeled VEGF association. We found a wide range of waiting times for tracks of any assembly state (Fig. 3F), indicating that many VEGF-VEGFR-2 association events occurred with pre-existing VEGFR-2 non-monomers. To investigate this issue further, we analyzed the relationship between VEGFR-2 merging events and VEGF association. Here also we found that, among the merging events that were positive for labeled VEGF, in half of them the association with VEGF occurred only after the merge of the two VEGFR-2 particles (Fig. 3G).

These analyses provide evidence that a good fraction of VEGF associates with pre-existing VEGFR-2 non-monomers, consistent with the alternative model of VEGF-VEGFR-2 binding and activation (Sarabipour et al., 2016). In light of the observation above that VEGF promotes interactions between imm/conf VEGFR-2 molecules, this suggests that the canonical and alternative models of VEGF-VEGFR-2 binding and activation co-exist in the native context, interestingly in a receptor mobility-dependent manner. Our data also suggest that many VEGF-VEGFR-2 binding and activation events follow an “in-between” model, where VEGF binds to a VEGFR-2 monomer, which then dimerizes with another VEGFR-2 monomer, but without any ligand-induced enhancement (Fig. 2F).

### VEGF stabilizes VEGFR-2 interactions

The simultaneous 2-color imaging experiments of VEGFR-2 and VEGF also allowed us to assess the effect of VEGF association on the stability of VEGFR-2 interactions, by comparing the dissociation rate of interaction events positive for VEGF to that of events negative for VEGF. Of note, because of labeling only 40% of VEGF molecules, some interaction events in the negative group could be bound to unlabeled VEGF, which would be invisible. Nevertheless, we found that the dissociation rate of interaction events positive for labeled VEGF was about half that of interaction events negative for labeled VEGF (Fig. 3H), indicating that association with VEGF stabilizes VEGFR-2 interactions. This stabilization most likely underlies the overall increase in VEGFR-2 assembly state upon VEGF addition (Fig. 3E) in spite of the decrease in overall merging rate (Fig. 2G).

### The fast decrease in D_mobile_ of VEGFR-2 upon VEGF stimulation is upstream of phosphorylation, while the shift toward the imm/conf mode is downstream

VEGF-VEGFR-2 binding leads to tyrosine phosphorylation of the intracellular domain of VEGFR-2. Thus, the observed effects of VEGF on VEGFR-2 behavior could be because of VEGF binding itself or because of VEGFR-2 phosphorylation and activation. To distinguish between these two scenarios, we performed single-molecule imaging followed by TCA for pHMVECs in the presence of the VEGFR phosphorylation inhibitor AAL-993 (Manley et al., 2002), using a 30 nM dose for 1 hour, which largely abolished VEGFR-2 activation by VEGF (Fig. S1C, D). These experiments also allowed us to investigate the role of VEGFR-2 phosphorylation in unstimulated cells, as previous studies have provided evidence for VEGFR-2 phosphorylation resulting from unligated dimerization (Sarabipour et al., 2016). While weak, we did in fact observe a slight reduction in VEGFR-2 phosphorylation in the presence of AAL-993 in unstimulated pHMVECs (Fig. S1C, D).

Diffusion analysis and TCA revealed that the two mobility modes of VEGFR-2 were conserved also in the presence of AAL-993, but now with a shift toward the mobile mode (i.e. opposite of the effect of VEGF), both in unstimulated and VEGF-stimulated cells (Fig. 4A-C). Parallel to the increase in mobile VEGFR-2 particles, there was a mild increase in the merging rate of VEGFR-2, both in the absence and presence of VEGF (Fig. S2). Interestingly, in contrast to the shift in modes from imm/conf to mobile, the almost immediate reduction of the diffusion coefficient of mobile VEGFR-2 particles (D_mobile_) upon VEGF stimulation was insensitive to the presence of AAL-993 (Fig. 4D).

**Figure 4.**
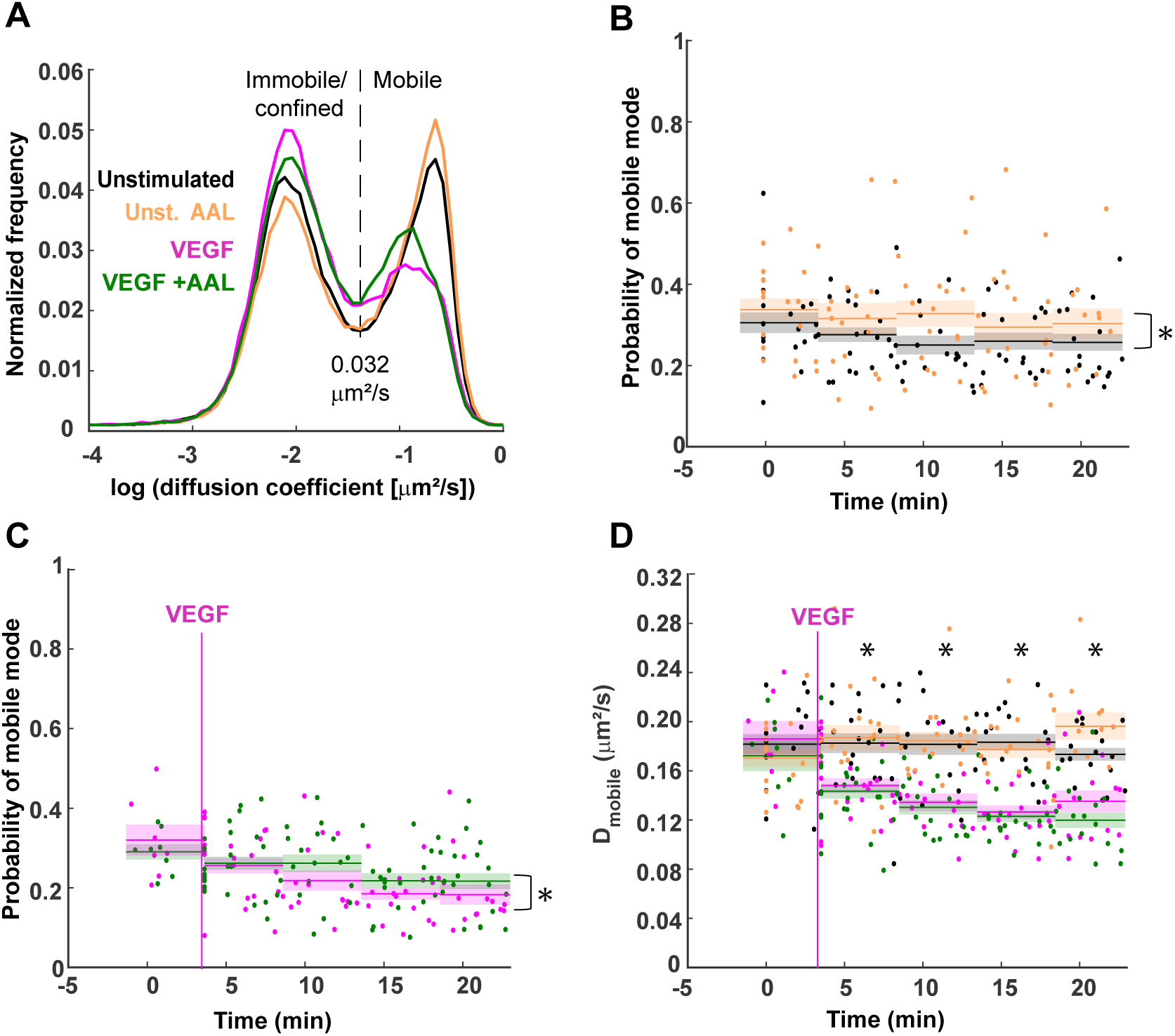
Effect of phosphorylation inhibition on the dynamic organization of VEGFR-2 on the pHMVEC surface. **(A)** VEGFR-2 diffusion coefficient distribution on the surface of pHMVECs: unstimulated (black), stimulated with 2 nM VEGF (magenta), unstimulated in the presence of 30 nM AAL-993 (orange), and VEGF-stimulated in the presence of AAL-993 (green). Unstimulated and VEGF-stimulated (both in the absence of AAL-993) distributions and vertical dashed line are a repeat of Fig. 2B. **(B-D)** Time-courses of the probability of VEGFR-2 molecules belonging to the mobile mode in unstimulated cells in the absence or presence of AAL-993 **(B)** and in VEGF-stimulated cells in the absence or presence of AAL-993 **(C)**, and the mean diffusion coefficient of mobile molecules in all conditions **(D)**. Dots, lines and surrounding shaded areas, and vertical magenta line as in Fig. 2C. Asterisks in (B-C): p-value < 0.02 for comparing the total time course between conditions using a paired t-test (in (C) the comparison is only for the time intervals after VEGF addition). Asterisks above time intervals in (D): p-value < 0.05 for comparing the measurements with and without VEGF stimulation in the presence of AAL per time interval using a t-test. N: Unstimulated: same as in Fig. 1; + VEGF: Same as in Fig. 2; unstimulated AAL: 133,815 tracks (48,851 with duration ≥ 5 frames) from 8 independent experiments with 7-9 cells each; AAL + VEGF: 134,371 tracks (54,198 with duration ≥ 5 frames) from 9 independent experiments with 7-8 cells each.

These observations indicate that VEGFR-2 phosphorylation upon VEGF binding plays a role in the shift in mobility mode from mobile to imm/conf, which in fact occurs on the timescale of the cellular signaling response (Fig. 2C and Fig. S1C, D). However, the almost immediate change in D_mobile_ upon VEGF binding is upstream of VEGFR-2 phosphorylation.

## Discussion

Through live-cell single-molecule imaging and a pipeline of automated image analysis and multiscale data analysis, we have exposed the previously unknown dynamic nanoscale organization of VEGFR-2 in its native plasma membrane on the EC surface and shed light on the interplay between this organization, ligand binding and receptor activation. We found that VEGFR-2 molecules on the surface of pHMVECs exhibit heterogeneity in both mobility and assembly state, where they can be mobile or imm/conf, both of which can be monomeric or non-monomeric, even in unstimulated cells (Fig. S3A-C). The mobile mode promotes VEGFR-2 encounters and interactions, which in turn enrich the imm/conf mode. The relationship however is not one-to-one: Many molecules remain mobile after interacting with each other, and, conversely, some imm/conf molecules appear to be monomeric. While the propensity for interactions depends on mobility mode, the dissociation rate of non-monomeric states is similar between the two modes.

Our work provides the first evidence that unligated VEGFR-2 molecules, expressed endogenously on the EC surface and with the full slew of native regulatory factors, can exist in non-monomeric states. Because of the sub-stoichiometric labeling inherent to single-molecule imaging, what we observe is most likely only the lower bound of the fraction of unligated VEGFR-2 molecules in non-monomeric states. Importantly, our analyses indicate that VEGF binds to both monomeric and non-monomeric VEGFR-2 (Fig. S3D-F), and provide evidence for the coexistence of the canonical and alternative models of VEGF-induced VEGFR-2 activation (Ruch et al., 2007; Sarabipour et al., 2016; Simons et al., 2016), and of an “in-between” model. In the “in-between” model, VEGF binds to a VEGFR-2 monomer (contrary to the alternative model), which then dimerizes with another VEGFR-2 monomer but without any ligand-induced enhancement (contrary to the canonical model), eventually leading to activation. The specific model followed by any particular VEGFR-2 molecule depends on its mobility mode and assembly state. In the particular case of molecules in a monomeric state, if they are in the imm/conf mode then they would follow the canonical model of VEGF-enhanced dimerization, while if they are in the mobile mode then they would follow the “in-between” model.

The partitioning of VEGFR-2 into two mobility modes is most likely due to many factors, and our work sheds light on these factors and their relationship to receptor activation. Specifically, we have found that VEGFR-2 phosphorylation promotes the imm/conf mode, even in unstimulated cells. Phosphorylation in the absence of stimulation could be due to unligated dimerization (Sarabipour et al., 2016) or association with Src family kinases (Chen et al., 1999; Jin et al., 2003; Tzima et al., 2005). Thus the imm/conf mode is at least partially due to interactions with downstream effectors and scaffolds (Labrecque et al., 2003; Lampugnani et al., 2006; Olsson et al., 2006), even if transiently in the absence of ligand. Interestingly, the time scale of the VEGF-induced shift from mobile to imm/conf mode parallels the time scale of VEGFR-2 activation by VEGF. In contrast, the diffusion coefficient of mobile VEGFR-2 molecules is independent of the phosphorylation state of VEGFR-2, and exhibits fast (almost immediate) change upon VEGF addition. Thus this change is most likely due to conformational changes in VEGFR-2 upon VEGF binding (Sarabipour et al., 2016) and/or the association of VEGFR-2 with co-receptors through VEGF (Gelfand et al., 2014; Somanath et al., 2009).

Our experiments and analyses demonstrate an intricate relationship between receptor mobility and interactions, leading to distinct effects of ligand and receptor activation steps based on the mobility of individual receptors. This highlights the need to study receptors in their native cellular environment in order to achieve a comprehensive understanding of receptor nanoscale organization, the factors that regulate it, and its relationship to receptor signaling. Live-cell single-molecule imaging is ideally suited for these purposes. Rigorous analytical approaches similar to (and building on) the approaches that we have developed here will be critical for making full use of the rich information contained in live-cell single-molecule imaging data in order to tease out the “rules” of molecular behavior and link them to their emergent cellular outcomes.

## Acknowledgments

We thank Dr. Tieqiao (Tim) Zhang for microscopy support, Dr. Ashley Lakoduk for sharing molecular biology expertise, and Drs. Sandra Schmid and Luke Rice for feedback on the manuscript. This work was supported by funding from NIH/NIGMS (R35 GM119619), The Welch Foundation (I-1901), CPRIT (R1216) and the UT Southwestern Endowed Scholars Program to KJ.

## Author contributions

B.R.A. and K.J. designed research; B.R.A., S.L. and A.D. performed experiments; A.R.V., L.R.O., T.K., M.K. and K.J. wrote analysis software; B.R.A. and K.J. performed analysis; B.R.A. and K.J. wrote the paper with input from all co-authors.

## STAR Methods

### Key resources table

| REAGENT or RESOURCE | SOURCE | IDENTIFIER |
| --- | --- | --- |
| **Antibodies** |  |  |
| Mouse monoclonal anti-VEGFR2 (EIC) | Abcam | Cat#ab9530 |
| Goat anti-mouse Fab secondary fragment (Rhodamine- Red-X conjugated) | Jackson ImmunoResearch | Cat#115-297-003<br>Lot#142110 |
| Rabbit monoclonal anti-VEGFR2 (55B11) | Cell Signaling Technology | Cat#2479 |
| Rabbit monoclonal anti-phospho VEGFR2 (Y1175) | Cell Signaling Technology | Cat#2478 |
| Mouse monoclonal anti-actin (AC40) | Millipore-Sigma | Cat#A4700 |
| Rabbit polyclonal anti-GAPDH | Abcam | Cat#ab9485 |
| Goat anti-Rabbit polyclonal antibody (HRP conjugated) | Thermo Scientific | Cat#G21234 |
| **Chemicals, Peptides, and Recombinant Proteins** |  |  |
| VEGF-A<sub>165</sub> (VEGF) | GenScript | Cat# Z03073-50 |
| AAL-993 | Millipore-Sigma | Cat#676504 |
| **Critical Commercial Assays** |  |  |
| Pierce Mouse IgG1 Fab and F(ab')<sub>2</sub> Preparation Kit | Thermo Scientific | Cat#44980 |
| Atto Protein Labeling Kit | Millipore-Sigma | Cat#38371 |
| **Experimental Models: Cell Lines** |  |  |
| Human: Dermal Microvascular Endothelial Cells- Neonatal (HD-MVECn), passages 4 to 7 | LifeLine Cell Technology | Cat#FC-0042 |
| **Software and Algorithms** |  |  |
| MATLAB | The MathWorks | N/A |
| Bio-formats | Linkert et al., 2010 | https://www.openmicroscopy.org/bio-formats/ |
| u-track | Jaqaman et al., 2008 | https://github.com/DanuserLab/u-track |
| Diffusion mode analysis | Jaqaman et al., 2016 | https://drive.google.com/open?id=1hp9jsLUJ2hT42QzKjtBDrpavDBZb7Tm9 (for the review process). Upon publication of this work, it will be made available via GitHub. |
| FISIK | De Oliveira and Jaqaman 2019 | https://github.com/kjaqaman/FISIK |
| Receptor-ligand coupled analysis | This manuscript | https://drive.google.com/open?id=1hp9jsLUJ2hT42QzKjtBDrpavDBZb7Tm9 (for the review process). Upon publication of this work, it will be made available via GitHub. |
| Time Course Analysis | This manuscript | https://drive.google.com/open?id=1hp9jsLUJ2hT42QzKjtBDrpavDBZb7Tm9 (for the review process). Upon publication of this work, it will be made available via GitHub. |

### Lead contact and materials availability

Further information and requests for resources should be directed to and will be fulfilled by the Lead Contact, Khuloud Jaqaman.

#### Materials availability statement

This study did not generate new unique reagents.

### Experimental model and subject details

This study employed early passage (P4-P7) primary cultures of human dermal microvascular endothelial cells isolated from neonatal foreskin (HD-MVECn, LifeLine Cell Technology, Frederick, MD). Cell type was authenticated by the supplier by ensuring that cells tested positive for Van Willebrand Factor (a marker for endothelial cells) and negative for smooth muscle α-actin. Cells were grown in VascuLife VEGF-Mv culture medium at 37°C + 5% CO2, and passaged every 48 hours upon reaching 70-90% confluency, following the supplier’s instructions.

### Method details

#### Cell culture and plating

Early passage (P4-P7) HD-MVECn cells (LifeLine Cell Technology, Frederick, MD) were grown in VascuLife VEGF-Mv culture medium for 48 h at 37°C + 5% CO2 until reaching 70-90% confluency. For stimulation, cells were treated with 2 nM VEGF-A_165_ (Genscript, Piscataway, NJ) at time points indicated in time-course analysis or Western Blots. To inhibit VEGFR-2 phosphorylation, cells were incubated with 30 nM of AAL-993 (MilliporeSigma, Burlington, MA), starting 1 h before the experiments and continuing for the duration of the experiments. For single molecule imaging experiments using TIRFM, 5.4 x 10^4^ cells were plated 18 h prior to imaging on base/acid cleaned glass bottom dishes (14 mm glass diameter with glass thickness of 1.5 mm, MatTek, Ashland, MA) pre-coated for 45 minutes with 10 µg/ml fibronectin (MilliporeSigma).

#### Preparation of anti-VEGFR-2 Fab fragments and fluorescence labeling of VEGF

The Pierce Mouse IgG1 Fab and F(ab’)2 Preparation Kit (Thermo Scientific, Waltham, MA) was used to generate Fab fragments from the anti-human VEGFR-2 mouse monoclonal antibody EIC (Abcam, Cambridge, MA). To reach optimal generation of Fab fragments, 500 µg of antibody were digested with immobilized ficin for 3h at 37°C, and, after the purification steps, the product was concentrated in a 10 kDa cut-off column. VEGF was conjugated with Atto488 through the succinimidyl ester group of the dye, using the Atto protein labeling kit following the manufacturer’s directions (MilliporeSigma). The dye-to-protein ratio obtained from the labeled VEGF was 0.4, as determined by spectrophotometry.

#### VEGFR-2 single molecule labeling

Plated cells were washed once with wash buffer (HBSS + 1 mM HEPES and 0.1% NGS), blocked for 15 min at 37°C in blocking buffer (1% BSA, 5% NGS in wash buffer) and incubated for 15 min with 0.2 µg/ml of primary anti-VEGFR-2 Fab fragments at 15°C. After 3 washes (1 min, RT), dishes were incubated for 15 min at RT in the dark with 2 µg/ml of a Rhodamine-Red-X (RRX)-conjugated goat anti-mouse secondary Fab fragment (Jackson ImmunoResearch, West Grove, PA). After 3 washes (1 min, RT), dishes were incubated with imaging buffer at 37°C (containing Oxyfluor 1%, Glucose 0.45%, Trolox 2nM) in order to reduce photobleaching. The purpose of the initial incubation with primary Fab fragments at 15°C, then incubation with secondary Fab fragments at RT, and then finally incubation with imaging buffer at 37°C was to minimize receptor and Fab fragment internalization during the labeling period while gently bringing the cells back to 37°C for imaging. VEGF or Atto488-VEGF to be added to cells were diluted in imaging buffer to a final concentration of 2 nM.

To determine the number of fluorophores per secondary fab fragment, the labeling procedure described above was followed, but with a much reduced concentration of primary Fab fragments (0.1 µg/ml) and of secondary Fab fragments (0.045 µg/ml).

#### Live-cell single-molecule imaging

Cells were imaged at 37°C using an S-TIRF system (Spectral Applied Research, via BioVision Technologies, Exton, PA) mounted on a Nikon Ti-E inverted microscope with Perfect Focus and a 60x/1.49 NA oil TIRF objective (Nikon Instruments, Melville, NY). The system was equipped with two Evolve EMCCD cameras (Photometrics, Tucson, AZ) for one-color or simultaneous two-color imaging, registered to within 1-3 pixels of each other. A custom 3x tube lens was employed to achieve an 89 nm pixel size in the recorded image. Illumination by a 561 nm diode pumped solid state laser (Cobolt) and/or a 488 nm diode laser (Coherent) was achieved through an ILE laser merge module (Spectral Applied Research), with 488 nm and 561 nm laser power of 3.65 mW and 13.2 mW, respectively, at the coverslip. The penetration depth was set to 80 nm via the Diskovery platform control (Spectral Applied Research). Videos were acquired with MetaMorph (Molecular Devices, San Jose, CA) in stream mode at 100 ms per frame (i.e. 10 Hz sampling) for 200 frames, using an EM gain of 100. Temperature and humidity were maintained using an environment chamber (Okolab, Otaviano, Italy), maintaining cell viability for the duration of the experiments. Every single-molecule imaging stream is preceded by a brightfield snapshot of the imaged cell region in order to delineate the cell mask (manually) for any ensuing analysis.

#### Western Blotting

After cell treatment (or not) with VEGF, Atto488-VEGF, and/or AAL-993 for the indicated times, cells were treated with lysis buffer (Mammalian-PE LB Buffer, GoldBio, St. Louis, MO) and a protease/phosphatase inhibitor cocktail (Thermo Scientific) for 10 min on ice under agitation. Lysates were collected after centrifugation, re-suspended in Laemmli sample buffer, and heated (90°C for 10 min). Samples were loaded in 4-15% gradient SDS-PAGE Mini-PROTEAN TGX gels (Bio-Rad, Hercules, CA). Proteins were transferred to PVDF membranes (100 V / 1 h / 4°C) and blocked using 5% BSA in TBST. Rabbit anti-human antibodies against VEGFR-2 (55B11, 1/2000 dilution, Cell Signaling, Danvers, MA), Y1175 phospho-VEGFR-2 (19A10, 1/1000 dilution, Cell Signaling), actin (1/1000 dilution, MilliporeSigma) and/or GAPDH (1/2500 dilution, Abcam) were incubated with membranes for 18h at 4°C. Goat anti-Rabbit secondary antibody conjugated with HRP (1/5000 dilution, Thermo Scientific) was incubated for 1 h at RT in the dark. Protein bands were detected by chemiluminescence using Clarity Western ECL Substrate (BioRad), and densitometric analyses were made using Image Lab (BioRad).

#### Computing environment

All image and data analysis tasks were performed in Matlab 2017a and more recent versions (The MathWorks, Natick, MA). Imaging streams were loaded into Matlab using Bio-Formats (Linkert et al., 2010).

#### Particle detection and tracking

Particle detection and tracking were performed using u-track (Jaqaman et al., 2008) (https://github.com/DanuserLab/u-track). The detection and tracking parameters, listed in Tables 1 and 2 below, were optimized based on the performance diagnostics included in u-track and visual inspection of the resulting detection and tracking. The mean particle localization precision, as estimated during the Gaussian mixture-model fitting step (Jaqaman et al., 2008), was 20 nm for VEGFR-2 in one-channel streams, and 26 nm and 33 nm for VEGFR-2 and VEGF, respectively, in two-channel streams. In all the following analyses, only detections and tracks inside the cell mask (determined from the brightfield image as described above) were employed.

**Table 1:** U-track detection parameters (Gaussian Mixture-Model Fitting). Shown are the parameters with non-default values for any of the three conditions, based on u-track Version 2.2.1.

| Detection parameters |  | One-channel imaging | Two-channel imaging |  |
| --- | --- | --- | --- | --- |
|  |  | VEGFR-2 | VEGFR-2 | VEGF |
| Gaussian standard deviation |  | 1.33 pixels |  |  |
| Local maxima detection | Use rolling window time-averaging | No | Yes (windows 1 and 3 frames)* |  |
| | α-value for comparison with local background | 0.05 | 0.05 (window 1), 0.1 (window 3) | 0.1 (both windows) |
| Gaussian mixture-model fitting at local maxima | Do iterative Gaussian mixture-model fitting | Yes |  |  |
| | α-value for residuals test | 0.01 | | 0.05 |
| | α-value for amplitude test | 0.05 | | 0.1 |
| | α-value for distance test | 0.01 | 0.05 | 0.1 |

\*A rolling window of 1 frame implies no time-averaging. A rolling window of 3 frames means averaging over the current frame, the frame before, and the frame after.

**Table 2:** U-track tracking parameters. Shown are the parameters with non-default values, based on u-track Version 2.2.1.

| Tracking parameters |  |  | VEGFR-2 from one- or two-channel imaging |
| --- | --- | --- | --- |
| Overall | Do segment merging |  | Yes |
|  | Do segment splitting |  | Yes |
| Frame-to-frame linking cost function | Brownian search radius upper bound |  | 10 pixels |
|  | Multiplication factor for Brownian search radius calculation |  | 4 |
| Gap closing, merging and splitting cost function | General | Brownian search radius upper bound | 10 pixels |
|  |  | Multiplication factor for Brownian search radius calculation | 4 |
|  |  | Scaling power to expand Brownian search radius | 0.25 |
|  |  | Use nearest neighbor distance to expand Brownian search radius | No |
|  | Merging & splitting | Search radius lower bound | 3 pixels |
|  |  | For an end/start, the possibility of merging/splitting is allowed only if there is no possibility of gap closing | Yes* |
|  | Birth & death | Birth and death cost | Derived from gap closing and merging/splitting cost formulae |

\*Limiting merging and splitting events to segment ends and starts that do not have the possibility of gap closing helps with distinguishing between specific interactions and particles passing by each other.

#### VEGFR-2 mobility analysis

The diffusion coefficient *D* of each VEGFR-2 particle with track duration ≥ 5 frames was calculated from its frame-to-frame displacement and localization precision, as previously described (Jaqaman et al., 2016). Tracks with duration < 5 frames were not used for mobility analysis. As described in the main text, *D* followed a bimodal distribution under all conditions (Fig. 5A). Thus, each VEGFR-2 track was classified as either mobile or immobile/confined, based on whether its diffusion coefficient was greater than or less than the classification threshold, respectively. The classification threshold was taken as 0.032 µm^2^/s for all conditions in one-channel streams (at the trough between the two modes). In the case of two-channel data, the distribution of VEGFR-2 diffusion coefficients was shifted slightly toward higher values, most likely because of the reduced localization precision in these data due to the images’ lower signal-to-noise ratio. Thus the classification threshold was shifted to 0.046 µm^2^/s in two-channel data, to coincide with the trough between the two modes in this condition.

#### VEGFR-2 assembly state and dissociation rate estimation

The apparent assembly state for the labeled subset of VEGFR-2 molecules was calculated from the particle intensities and their sequence of merging and splitting events as described in (De Oliveira and Jaqaman, 2019) (https://github.com/kjaqaman/FISIK). This analysis required the mean intensity of an individual fluorophore, which was estimated per imaging stream by decomposing the distribution of detected particle intensities in the first 5 frames of the stream into a superposition of multiple modes corresponding to 1, 2, 3, etc. fluorophores (Jaqaman et al., 2011). Each mode was taken as a log-normal distribution, where the mean and standard deviation of mode n were (approximately) n times those of mode 1 (Mutch et al., 2007). The fit was achieved using least squares, and the number of modes was determined using the Bayesian Information Criterion for model selection (Jaqaman and Danuser, 2006). With this, the mean individual fluorophore intensity was obtained per imaging stream, and then used to estimate the assembly state of each particle as described in (De Oliveira and Jaqaman, 2019). The dissociation rate per assembly state was then calculated from the transitions between assembly states as described in (De Oliveira and Jaqaman, 2019).

#### Analysis of VEGF-VEGFR-2 association in two-channel imaging streams

VEGF-VEGFR-2 association was determined in two steps:

First, in each frame, the colocalization between VEGFR-2 and VEGF particles was determined by solving a linear assignment problem (Jonker and Volgenant, 1987) based on their relative positions, with a maximum allowed distance of 4 pixels (∼350 nm). This maximum distance accounted for both the particles’ localization precision (0.3-0.4 pixels on average) and the 1-3 pixel registration shift between the two cameras acquiring the two-channel images. After this first step, each VEGFR-2 track had a 1/0 association flag with VEGF per frame.

Second, the association history per track was used to distinguish spurious associations from reliable ones. Specifically, a VEGFR-2 track was considered to reliably associate with VEGF if the association history satisfied the following two conditions: (i) Its association flag = 1 in at least three frames. (ii) Its association flag = 1 for at least 10% of the frames between its first frame of VEGF association and last frame of VEGF association (e.g., if the first frame of VEGF association is frame 11, and the last frame of VEGF association is frame 50, then the association flag should be 1 for at least 0.1 × 40 = 4 frames). For tracks considered to reliably associate with VEGF, the average fraction of time associated with VEGF was 57% of their duration (34% of the tracks were associated with VEGF for 90% of their duration).

With this, each VEGFR-2 track reliably associated with VEGF was divided into the interval before VEGF association and the interval during VEGF association for further analysis as needed (e.g. VEGFR-2 mobility). Note that if the VEGF-VEGFR-2 association was lost in some later frame, a VEGFR-2 track would then also have an interval after VEGF association, but this was not used for any analysis in our study.

#### Time-course analysis: data aggregation strategies

For time course analysis, two data aggregation strategies were taken, based on the amount of data available per individual cell to calculate the single-molecule property of interest.

Strategy 1. For single-molecule properties with sufficient data points per cell, primarily properties related to receptor mobility, single-molecule properties were calculated per cell (e.g. mean D_mobile_ per cell), and then individual cell measurements were grouped by time interval for time course analysis. A time interval of 5 minutes was used, as it provided enough individual cell measurements per time interval (10-20) to minimize measurement noise, while allowing for sufficient temporal sampling to capture temporal trends. In this case, the figures display the individual cell measurements, and the mean value and standard error of the mean for the grouped cells per time interval (e.g. Fig. 3C).

Strategy 2. For single-molecule properties with insufficient data points per cell, particularly properties related to receptor interactions (as seen in Fig. 3F, the rate of merging is on the order to 10^−5^/s/particle), the individual cell measurements could exhibit relatively large fluctuations from cell to cell. In this case, the grouping was done at the level of the tracks within the cells of interest. Specifically, the single-molecule property (e.g. dissociation rate) was calculated from all the tracks combined, and the standard deviation of that property was calculated via bootstrapping (using 100 bootstrap samples). This standard deviation was equivalent to the standard error of the mean of individual cell measurements. In this case, the figures display the property as calculated from the combined tracks and the standard deviation from bootstrapping (e.g. Fig. 2C or Fig. 4B).

Note that, when such analysis was performed on unstimulated cells, all cells were grouped together, as VEGFR-2 properties did not vary over time in that case (e.g. Fig. 3C and Fig. 4E). On the other hand, when such analysis was performed on stimulated cells, a time interval of 10 minutes was used, thus separating the early and late trends in VEGFR-2 behavior upon VEGF addition (0-10 min and 10-20 min after VEGF addition, respectively). A 10-minute time interval was used instead of a 5-minute time interval (as done above) in order to further reduce measurement noise.

### Quantification and statistical analysis

#### Dataset information

The statistical details of all experiments (number of independent repeats, number of cells imaged per repeat, and number of tracks used for analysis) are reported in the figure legends. Statistical tests are described in the figure legends, and in more detail below.

#### Statistical tests

Below is a description of the various statistical tests used for comparing single-molecule properties:

For properties where individual cell measurements were available in the time course analysis (data aggregation strategy 1, described in the previous section), the properties were compared between conditions for each time interval using a one-sided t-test (based on the individual cell measurements within the time interval). In this case, significant differences are indicated by asterisks above each time interval (e.g. Fig. 3C).

In cases where all per-interval comparisons were not significant (p-value > 0.05), the full time course was compared between conditions using a paired, one-sided t-test. Specifically, each time course was represented by its series of mean values per time interval, and the series of two conditions were compared using a paired t-test. In other words, time courses without VEGF stimulation had 5 data points (e.g. Fig. 5B), while time courses with VEGF stimulation had 4 data points (corresponding to the time intervals after VEGF addition; e.g. Fig. 5C). In this case, significant differences are indicated by a square bracket and an asterisk at the end of the time course plot (e.g. Fig. 5B). The reasoning behind this test was that temporal persistence in a trend (e.g. one condition being always higher than the other) might indicate a significant difference, even if the difference is not strong enough to detect on a per time interval basis.

For properties calculated by first combining all tracks for a group of cells (data aggregation strategy 2, described in the previous section), the property standard deviation was obtained via bootstrapping, and was equivalent to the standard error of the mean of individual cell measurements. In this case, to compare conditions (or time intervals) 1 and 2, with property values µ_1_ and µ_2_, and variances v_1_ and v_2_, the difference between them, µ_1_ – µ_2_, was taken to follow *N*(0, v_1_+v_2_). This distribution was then used to calculate the p-value to assess the difference between the two conditions (or time intervals). In this case, significant differences are indicated by asterisks above each time interval in the time course (e.g. Fig. 4C), or otherwise as described in the figure legends.

### Data and code availability

The data generated during this study are available from the corresponding author upon request. All custom code written for the purposes of this study will be made available via GitHub (under https://github.com/kjaqaman) upon manuscript publication.

## Supplemental Material

**Figure S1.**
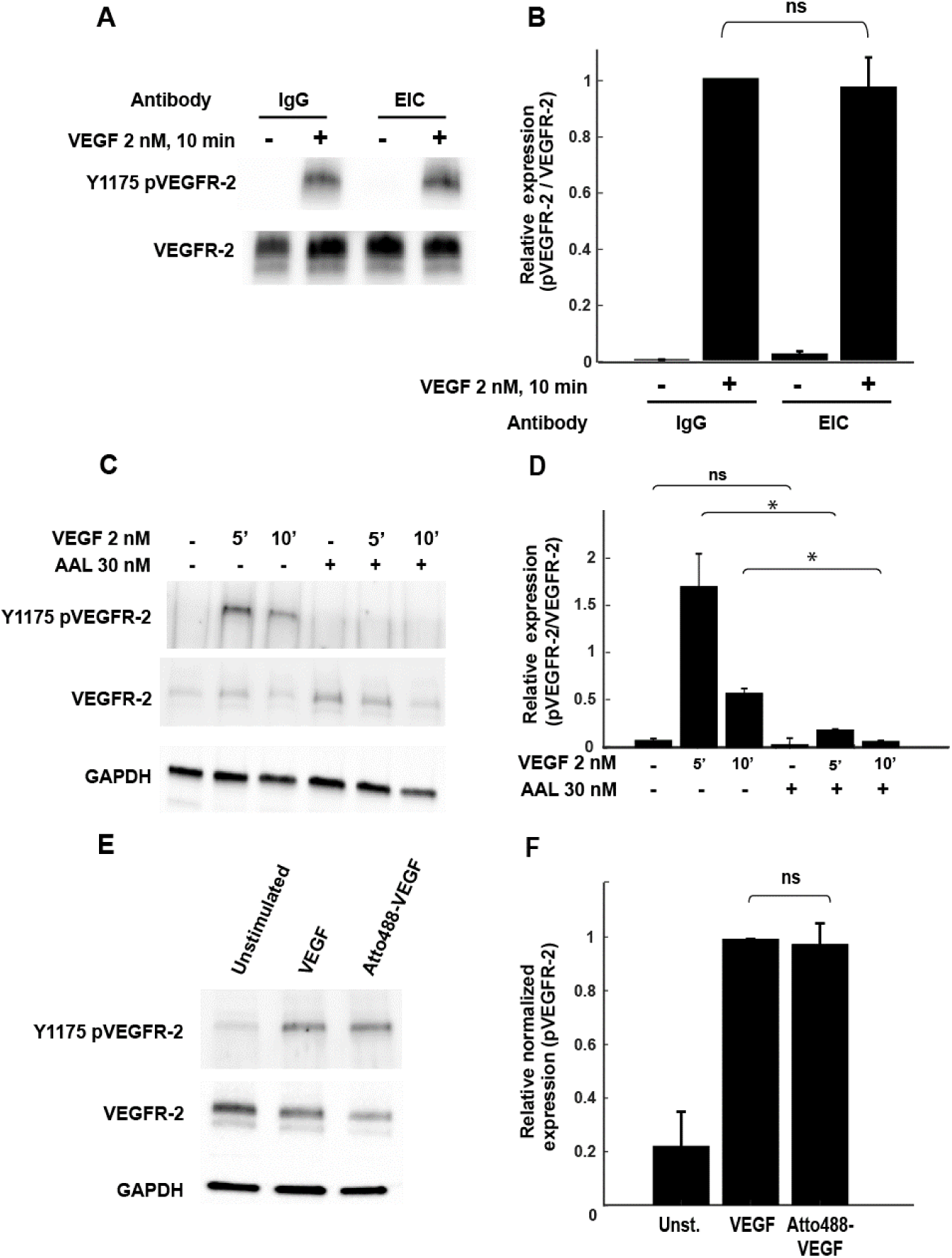
Validation of reagents. **(A-B)** Assessment of EIC antibody neutrality. **(A)** Representative Western blot from HMVECs incubated for 4 h in the presence of 5 µg/ml of either nonspecific IgG1 or the anti-VEGFR-2 monoclonal antibody EIC, and stimulated or not with 2 nM VEGF for 10 minutes. Expression of phosphorylated VEGFR-2 (Y1175 pVEGFR-2) and total VEGFR-2 were detected, as described in Materials and Methods. **(B)** Phosphorylated VEGFR-2 expression relative to total VEGFR-2 expression, measured by densitometry, from N = 4 Western blot repeats. Error bars: standard error of the mean. No significant difference was observed between cells treated with IgG1 or EIC antibodies (ns: not significant (p > 0.05, Student’s t-test)). **(C-D)** Time course of VEGFR-2 phosphorylation in the presence or absence of VEGF and/or AAL-993. **(C)** Representative Western blot from HMVECs stimulated or not with 2 nM VEGF for 5 or 10 minutes, in the presence or not of the VEGFR-2 phosphorylation inhibitor AAL-993 (treatment with 30 nM for 1 h). Expression of phosphorylated VEGFR-2 (Y1175 pVEGFR-2), total VEGFR-2, and GAPDH were detected, as described in Materials and Methods. **(D)** Phosphorylated VEGFR-2 expression relative to total VEGFR-2 expression, measured by densitometry, from N = 3 Western blot repeats. Error bars: standard error of the mean. Asterisks and ns (not significant): p-value < 0.05 and p-value > 0.05, respectively (Student’s t-test). As expected, VEGF leads to significant VEGFR-2 phosphorylation within 5 min, and this is largely abolished in the presence of AAL-993. **(E-F)** Validation of Atto488-VEGF. **(E)** Representative Western blot from HMVECs stimulated or not with 2nM VEGF or 2 nM Atto488-VEGF (of which 40% is actually labeled) for 5 minutes. Expression of phosphorylated VEGFR-2 (Y1175 pVEGFR-2), total VEGFR-2, and GAPDH were detected, as described in Materials and Methods. **(F)** Relative normalized expression of phosphorylated VEGFR-2, measured by densitometry, from N = 3 Western blot repeats. Error bars: standard error of the mean. No significant difference was observed between cells treated with VEGF or Atto488-VEGF (ns: not significant (p > 0.05, Student’s t-test)).

**Figure S2.**
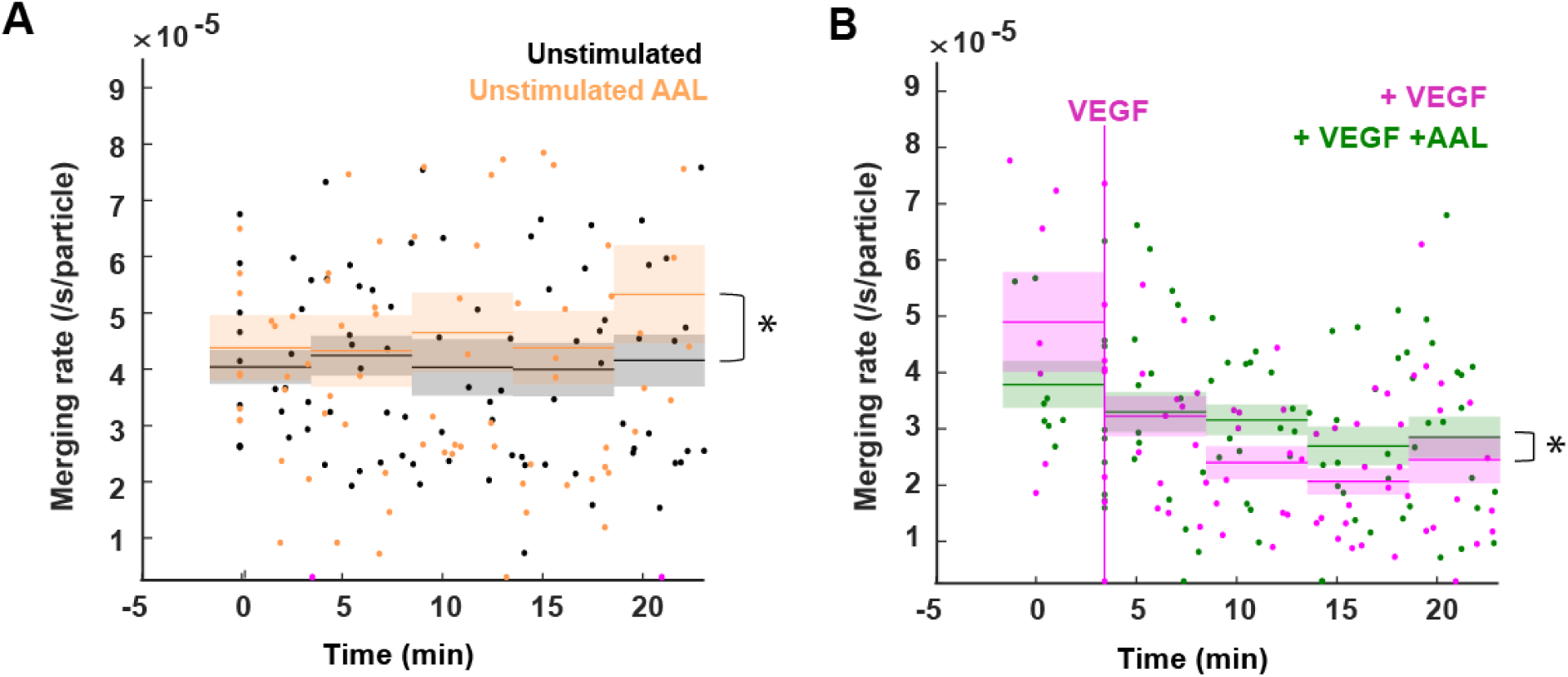
Effect of phosphorylation inhibition on the merging rate of VEGFR-2. **(A-B)** Time courses of the merging rates in unstimulated cells in the absence (black) or presence (orange) of 30 nM AAL-993 **(A)**, or in cells stimulated with 2 nM VEGF in the absence (magenta) or presence (green) of 30 nM AAL-993 **(B)**. Dots, lines and surrounding shaded areas, and vertical magenta line as in Fig. 2C. Asterisks: p-value < 0.03 for comparing the total time course between conditions using a paired t-test (in (B) the comparison is only for the time intervals after VEGF addition). N as in Fig. 4.

**Figure S3.**
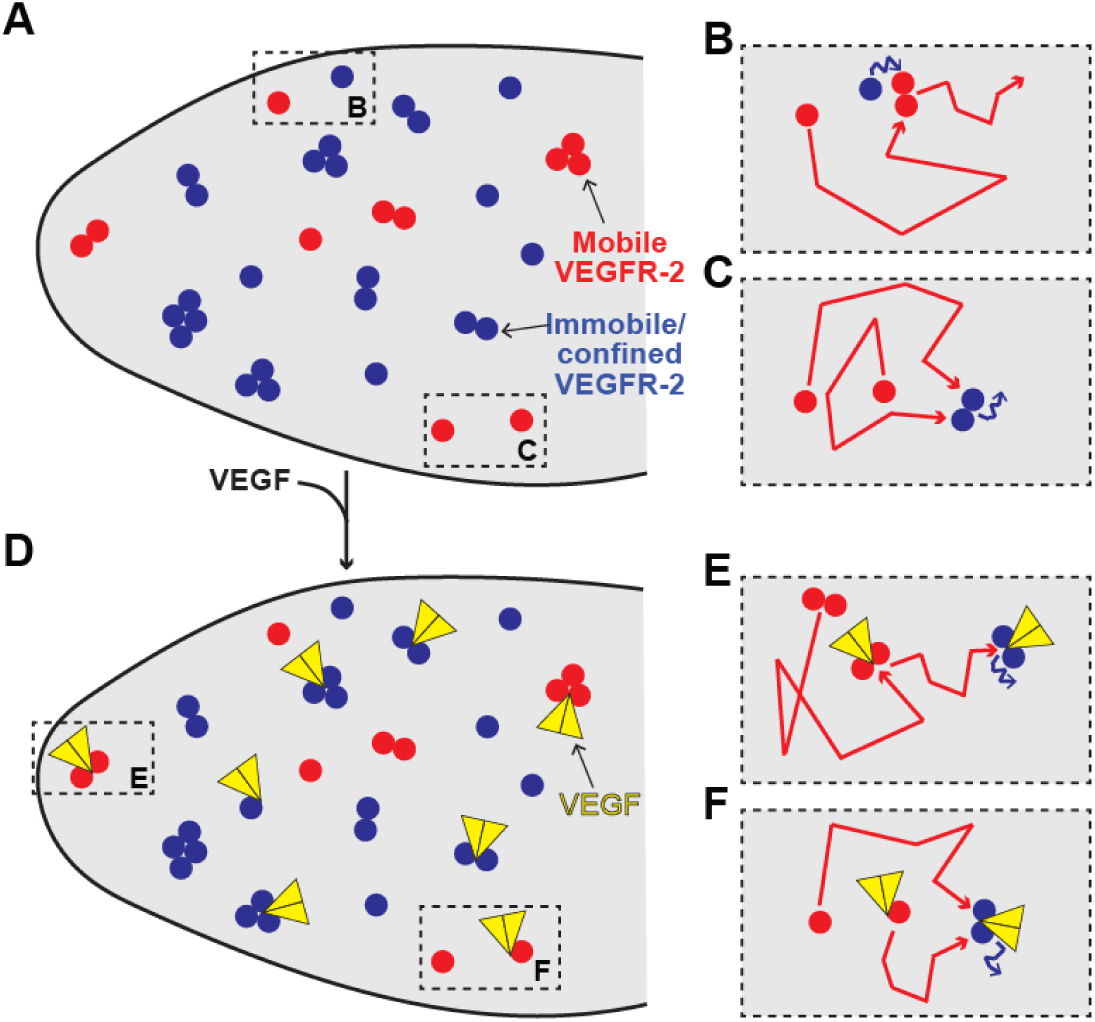
Schematic of VEGFR-2 spatiotemporal organization and its relationship to VEGF in pHMVECs. **(A)** In unstimulated cells, VEGFR-2 can be mobile (red; ∼35%) or imm/conf (blue; ∼65%), and has a distribution of assembly states, with more imm/conf molecules than mobile molecules in non-monomeric states. Boxed regions are used to illustrate examples of the interplay between mobility and interactions in (B-C). **(B-C)** Example illustrations of one mobile monomer and one imm/conf monomer associating to produce a mobile dimer **(B)**, and of two mobile monomers associating to produce an imm/conf dimer **(C)**. The examples illustrate a sequence of events, showing the molecules when separate (separate dots) and upon interacting with each other (paired-dots). The dot colors reflect the molecules’ mobility mode moving forward. Molecule movement is indicated by jagged lines colored by mobility mode, with arrows indicating direction of movement. While these two examples are not comprehensive, they reflect that the mobile mode promotes interactions, which in turn enrich the imm/conf mode, but without a simple one-to-one relationship between mobility and assembly state. (D) VEGF binds to VEGFR-2 monomers and pre-existing non-monomers. VEGF symbol is not drawn to scale. Boxed regions are used to illustrate examples of the interplay between VEGF binding and VEGFR-2 mobility and interactions (E-F). **(E-F)** Example illustrations of VEGF binding to a pre-existing mobile dimer, such that the ligated dimer slows down and eventually switches mobility to imm/conf **(E)**, and of VEGF binding to a mobile monomer, which then associates with an unligated mobile monomer, producing a ligated imm/conf dimer. The illustration style follows that in (B-C). Again, while these two examples are not comprehensive, they reflect that VEGF can bind monomeric and non-monomeric VEGFR-2 and that VEGF binding slows down mobile VEGFR-2 and promotes the imm/conf mode of VEGFR-2 mobility, which depends on VEGFR-2 phosphorylation (not illustrated).

## Movie legends

**Movie 1: Single-molecule imaging of endogenous VEGFR-2 in pHMVECs.** Representative 10 Hz single-molecule imaging stream of an unstimulated cell. The first 10 s of the 20 s stream are shown. Image size is 45.5 × 45.5 µm^2^.

**Movie 2: VEGFR-2 particle tracking.** Same single-molecule imaging stream as in Movie 1, with VEGFR-2 tracks overlaid on the fluorescence images. Random track colors are used to help distinguish neighboring tracks.

**Movie 3: VEGFR-2 mobility classification.** Same VEGFR-2 tracks as in Movie 2, but now color-coded based on mobility mode. Red = mobile; blue = immobile/confined, light pink = too short to classify (duration < 5 frames).

**Movie 4: Example VEGFR-2 merging and splitting events.** Example is a zoom of an area from same stream shown in Movies 1-3. Image size is 4 × 4.8 µm^2^.

**Movie 5: Example of a VEGFR-2 particle associated with a VEGF particle.** Example is from a 10 Hz/20 s simultaneous two-channel TIRF imaging stream of VEGFR-2 labeled with RRX (left image) and Atto488-VEGF (right image). The VEGFR-2 track is overlaid on both images as a red circle followed by a light pink tail, while the associated VEGF particle is overlaid on the right image as a cyan circle. Image size is 3.2 × 3.2 µm^2^.

## References

Bogdanovic, E., Coombs, N., and Dumont, D.J. (2009). Oligomerized Tie2 localizes to clathrin-coated pits in response to angiopoietin-1. Histochem Cell Biol 132, 225–237.

Casaletto, J.B., and McClatchey, A.I. (2012). Spatial regulation of receptor tyrosine kinases in development and cancer. Nat Rev Cancer 12, 386–399.

Chen, K.D., Li, Y.S., Kim, M., Li, S., Yuan, S., Chien, S., and Shyy, J.Y.J. (1999). Mechanotransduction in response to shear stress - Roles of receptor tyrosine kinases, integrins, and Shc. J Biol Chem 274, 18393–18400.

Chung, I., Akita, R., Vandlen, R., Toomre, D., Schlessinger, J., and Mellman, I. (2010). Spatial control of EGF receptor activation by reversible dimerization on living cells. Nature 464, 783–787.

De Oliveira, L.R., and Jaqaman, K. (2019). FISIK: Framework for the Inference of In Situ Interaction Kinetics from Single-Molecule Imaging Data. Biophys J 117, 1012–1028.

Ferrara, N., Gerber, H.P., and LeCouter, J. (2003). The biology of VEGF and its receptors. Nat Med 9, 669–676.

Freeman, S.A., Vega, A., Riedl, M., Collins, R.F., Ostrowski, P.P., Woods, E.C., Bertozzi, C.R., Tammi, M.I., Lidke, D.S., Johnson, P., et al. (2018). Transmembrane Pickets Connect Cyto- and Pericellular Skeletons Forming Barriers to Receptor Engagement. Cell 172, 305–317 e310.

Fuh, G., Li, B., Crowley, C., Cunningham, B., and Wells, J.A. (1998). Requirements for binding and signaling of the kinase domain receptor for vascular endothelial growth factor. J Biol Chem 273, 11197–11204.

Garcia-Parajo, M.F., Cambi, A., Torreno-Pina, J.A., Thompson, N., and Jacobson, K. (2014). Nanoclustering as a dominant feature of plasma membrane organization. J Cell Sci 127, 4995–5005.

Gelfand, M.V., Hagan, N., Tata, A., Oh, W.J., Lacoste, B., Kang, K.T., Kopycinska, J., Bischoff, J., Wang, J.H., and Gu, C. (2014). Neuropilin-1 functions as a VEGFR2 co-receptor to guide developmental angiogenesis independent of ligand binding. Elife 3, e03720.

Githaka, J.M., Vega, A.R., Baird, M.A., Davidson, M.W., Jaqaman, K., and Touret, N. (2016). Ligand-induced growth and compaction of CD36 nanoclusters enriched in Fyn induces Fyn signaling. J Cell Sci 129, 4175–4189.

Hamdollah Zadeh, M.A., Glass, C.A., Magnussen, A., Hancox, J.C., and Bates, D.O. (2008). VEGF-mediated elevated intracellular calcium and angiogenesis in human microvascular endothelial cells in vitro are inhibited by dominant negative TRPC6. Microcirculation (New York, NY : 1994) 15, 605–614.

Jaqaman, K., and Danuser, G. (2006). Linking data to models: data regression. Nat Rev Mol Cell Biol 7, 813–819.

Jaqaman, K., Galbraith, J.A., Davidson, M.W., and Galbraith, C.G. (2016). Changes in single-molecule integrin dynamics linked to local cellular behavior. Mol Biol Cell 27, 1561–1569.

Jaqaman, K., Kuwata, H., Touret, N., Collins, R., Trimble, W.S., Danuser, G., and Grinstein, S. (2011). Cytoskeletal control of CD36 diffusion promotes its receptor and signaling function. Cell 146, 593–606.

Jaqaman, K., Loerke, D., Mettlen, M., Kuwata, H., Grinstein, S., Schmid, S.L., and Danuser, G. (2008). Robust single-particle tracking in live-cell time-lapse sequences. Nat Methods 5, 695–702.

Jin, Z.G., Ueba, H., Tanimoto, T., Lungu, A.O., Frame, M.D., and Berk, B.C. (2003). Ligand-independent activation of vascular endothelial growth factor receptor 2 by fluid shear stress regulates activation of endothelial nitric oxide synthase. Circ Res 93, 354–363.

Jonker, R., and Volgenant, A. (1987). A Shortest Augmenting Path Algorithm for Dense and Sparse Linear Assignment Problems. Computing 38, 325–340.

Karaman, S., Leppanen, V.M., and Alitalo, K. (2018). Vascular endothelial growth factor signaling in development and disease. Development 145, 8.

King, C., and Hristova, K. (2019). Direct measurements of VEGF-VEGFR2 binding affinities reveal the coupling between ligand binding and receptor dimerization. J Biol Chem 294, 9064–9075.

Labrecque, L., Royal, I., Surprenant, D.S., Patterson, C., Gingras, D., and Beliveau, R. (2003). Regulation of vascular endothelial growth factor receptor-2 activity by caveolin-1 and plasma membrane cholesterol. Mol Biol Cell 14, 334–347.

Lampugnani, M.G., Orsenigo, F., Gagliani, M.C., Tacchetti, C., and Dejana, E. (2006). Vascular endothelial cadherin controls VEGFR-2 internalization and signaling from intracellular compartments. J Cell Biol 174, 593–604.

Lin, C.-C., Melo, Fernando A., Ghosh, R., Suen, Kin M., Stagg, Loren J., Kirkpatrick, J., Arold, Stefan T., Ahmed, Z., and Ladbury, John E. (2012). Inhibition of Basal FGF Receptor Signaling by Dimeric Grb2. Cell 149, 1514–1524.

Linkert, M., Rueden, C.T., Allan, C., Burel, J.M., Moore, W., Patterson, A., Loranger, B., Moore, J., Neves, C., MacDonald, D., et al. (2010). Metadata matters: access to image data in the real world. J Cell Biol 189, 777–782.

Liu, Z., Lavis, L.D., and Betzig, E. (2015). Imaging live-cell dynamics and structure at the single-molecule level. Mol Cell 58, 644–659.

Low-Nam, S.T., Lidke, K.A., Cutler, P.J., Roovers, R.C., van Bergen en Henegouwen, P.M., Wilson, B.S., and Lidke, D.S. (2011). ErbB1 dimerization is promoted by domain co-confinement and stabilized by ligand binding. Nat Struct Mol Biol 18, 1244–1249.

Manley, P.W., Furet, P., Bold, G., Bruggen, J., Mestan, J., Meyer, T., Schnell, C.R., Wood, J., Haberey, M., Huth, A., et al. (2002). Anthranilic acid amides: a novel class of antiangiogenic VEGF receptor kinase inhibitors. J Med Chem 45, 5687–5693.

Mischel, P.S., Umbach, J.A., Eskandari, S., Smith, S.G., Gundersen, C.B., and Zampighi, G.A. (2002). Nerve Growth Factor Signals via Preexisting TrkA Receptor Oligomers. Biophysical Journal 83, 968–976.

Mutch, S.A., Fujimoto, B.S., Kuyper, C.L., Kuo, J.S., Bajjalieh, S.M., and Chiu, D.T. (2007). Deconvolving single-molecule intensity distributions for quantitative microscopy measurements. Biophysical Journal 92, 2926–2943.

Olsson, A.K., Dimberg, A., Kreuger, J., and Claesson-Welsh, L. (2006). VEGF receptor signalling - in control of vascular function. Nat Rev Mol Cell Biol 7, 359–371.

Ruch, C., Skiniotis, G., Steinmetz, M.O., Walz, T., and Ballmer-Hofer, K. (2007). Structure of a VEGF–VEGF receptor complex determined by electron microscopy. Nature Structural & Molecular Biology 14, 249.

Sarabipour, S., Ballmer-Hofer, K., and Hristova, K. (2016). VEGFR-2 conformational switch in response to ligand binding. Elife 5, e13876.

Simons, M., Gordon, E., and Claesson-Welsh, L. (2016). Mechanisms and regulation of endothelial VEGF receptor signalling. Nat Rev Mol Cell Biol 17, 611–625.

Somanath, P.R., Malinin, N.L., and Byzova, T.V. (2009). Cooperation between integrin alphavbeta3 and VEGFR2 in angiogenesis. Angiogenesis 12, 177–185.

Sungkaworn, T., Jobin, M.L., Burnecki, K., Weron, A., Lohse, M.J., and Calebiro, D. (2017). Single-molecule imaging reveals receptor-G protein interactions at cell surface hot spots. Nature 550, 543–547.

Terman, B.I., Dougher-Vermazen, M., Carrion, M.E., Dimitrov, D., Armellino, D.C., Gospodarowicz, D., and Bohlen, P. (1992). Identification of the KDR tyrosine kinase as a receptor for vascular endothelial cell growth factor. Biochem Biophys Res Commun 187, 1579–1586.

Treanor, B., Depoil, D., Gonzalez-Granja, A., Barral, P., Weber, M., Dushek, O., Bruckbauer, A., and Batista, F.D. (2010). The Membrane Skeleton Controls Diffusion Dynamics and Signaling through the B Cell Receptor. Immunity 32, 187–199.

Tzima, E., Irani-Tehrani, M., Kiosses, W.B., Dejana, E., Schultz, D.A., Engelhardt, B., Cao, G., DeLisser, H., and Schwartz, M.A. (2005). A mechanosensory complex that mediates the endothelial cell response to fluid shear stress. Nature 437, 426–431.

Xia, T., Li, N., and Fang, X. (2013). Single-molecule fluorescence imaging in living cells. Annu Rev Phys Chem 64, 459–480.

Zhou, Y., Prakash, P., Liang, H., Cho, K.J., Gorfe, A.A., and Hancock, J.F. (2017). Lipid-Sorting Specificity Encoded in K-Ras Membrane Anchor Regulates Signal Output. Cell 168, 239-+.

